# Fishing pressure induces changes in DNA methylation in genetically homogeneous marine metapopulations

**DOI:** 10.64898/2026.03.19.712898

**Authors:** M. Barcelo-Serra, A.C. Mateman, A.S. Pijl, J.E. Risse, B. Sepers, M.A. Cortes-Pujol, J. Alós, K. van Oers

## Abstract

Trait-selective harvesting by fisheries can impose strong selective pressures on fish populations, driving changes in life history traits affecting fisheries productivity and ecosystem functioning. While the genetic consequences of harvesting have been extensively studied, the extent to which phenotypic variation reflects genomic evolution versus environmentally-induced plasticity remains unclear. Epigenetic mechanisms, such as DNA methylation, may mediate between these processes, serving as a rapid and reversible response to the selective pressures imposed by harvesting. In this study, we implemented an improved laboratory and bioinformatics protocol, epiGBS3, to examine genomic variation and DNA methylation patterns in the marine fish *Xyrichtys novacula*. The study spanned three replicated geographical areas each comprising two adjacent locations: an intensively exploited fishery and a no-take Marine Protected Area (ntMPA). A nested analysis design across the three areas revealed strong gene flow and no evidence of genetic structure. Nevertheless, nucleotide diversity was significantly reduced in fisheries relative to ntMPAs. We also found that DNA methylation levels differed between protected and exploited sites after controlling for age, suggesting that fishing may influence epigenetic changes independently of fisheries-induced age-truncation effects. This represents one of the first lines of evidence that fisheries can potentially shape epigenetic variation, supporting DNA methylation as contributor to local adaptation under high gene flow and strong anthropogenic selection.

## Introduction

Human harvesting of wild fish populations has intensified over the past centuries, driven by industrialization and increased demand of marine-derived products [1]. In the early 20^th^ century, researchers witnessing this intensification proposed fishing as a potential agent of artificial selection capable of driving phenotypic change in exploited populations [2,3]. Since then, numerous studies have investigated the effects of trait-selective harvesting on fish biology and its potential evolutionary consequences on fish stocks (reviewed in [4]). Despite this extensive body of work, debate remains over the extent and mechanisms of fisheries-induced evolution, specifically whether observed phenotypic changes arise from heritable genetic shifts or environmentally induced plasticity [5,6]. While this debate persists, recent studies increasingly provide genomic evidence consistent with fisheries-induced evolution. These advances, largely enabled by the advances in genomic tools, allow for the detection of polygenic responses involving many loci with small individual effects [7,8].

Evidence from artificial selection experiments, simulation models and studies on exploited wild populations demonstrates that both commercial and recreational fishing can impose strong directional selection on morphological, physiological and behavioural traits [9,10]. Selective capture of larger, bolder and more catchable individuals, along with indirect selection on correlated traits, has been associated with reduced body size, earlier maturation and behavioural shifts such as increased timidity [11–14]. These trait changes can have profound demographic and ecological consequences. Trait-selective harvesting can reduce reproductive output, disrupt courtship dynamics and alter mate choice, ultimately lowering reproductive success and increasing the risk of population decline and collapse [10,14]. Although artificial selection experiments confirm that fishing can induce evolutionary responses, and recent genomic studies increasingly report genetic differentiation associated with harvesting pressure [7,8], phenotypic divergence in many exploited wild populations often occur rapidly, on ecological timescales that are hard to explain based on genetic change alone [15]. This suggests that additional mechanisms may contribute to fast and potentially reversible phenotypic responses to harvesting pressure.

Epigenetic modifications such as DNA methylation (the covalent addition of a methyl group to cytosines in CpG dinucleotides) can facilitate plastic phenotypic responses in vertebrates without altering the underlying DNA sequence [16–18]. DNA methylation can modulate gene expression, transcriptional activity, alternative splicing, transcriptional noise, enhancer regulation and alternative promoter usage, acting on diverse genomic features including promoters, exons and introns [19–23]. Furthermore, changes in DNA methylation can be environmentally induced and reversible, providing an additional layer of molecular flexibility under rapid environmental change [16].

In the context of fishing-induced changes in population density, demography, sex ratio and social structure, DNA methylation may enable rapid, plastic adjustments in key life-history traits. For example, in sequential hermaphrodites, the selective removal of one sex can alter the timing of sex transition [24] and such shifts could be mediated by DNA methylation changes that accelerate sex transition and promote earlier maturation. Likewise, the disruption of social hierarchies, affecting dominance and access to resources, could affect behavioural and growth patterns [25] through comparable epigenetic mechanisms. These fishery-induced environmental effects could potentially alter DNA methylation patterns in surviving individuals, yet no studies have evaluated whether such effects contribute to the phenotypic variation observed in exploited populations.

No-take Marine Protected Areas (ntMPAs) have been proposed as management tools to buffer the potential molecular effects of trait-selective harvesting by providing spatial refuges for exploited species and protecting ecosystems as a whole [26,27]. In fact, ntMPAs have been shown to restore body size distributions and natural sex ratios [28]. Many exploited marine species exhibit pelagic early-life stages and high dispersal potential, leading to strong hydrodynamic connectivity across habitats. As a result, ntMPAs hydrodynamically connected to fisheries, can compensate for the loss imposed by selective harvesting by migration and gene flow [27]. Understanding how fishing shapes both genetic and DNA methylation variation is critical for assessing population resilience and the role of ntMPAs as molecular reservoirs.

In this study, we investigate the impact of fishing pressure on both genomic and epigenetic patterns (DNA methylation) in the coastal marine fish pearly razorfish (*Xyrichtys novacula*; Suppl. Fig. S1). Using a replicated study design considering ntMPAs adjacent to intensively exploited fisheries as natural contrasts in selection regimes, we test the hypothesis that trait-selective harvesting alters genetic and DNA methylation variation. Specifically, we examine whether fishing reduces genetic diversity through selective removal of phenotypes, whether compensatory gene flow from ntMPAs mitigates genetic erosion and whether DNA methylation profiles diverge between protection levels in ways consistent with environmental induced plasticity. By integrating genetic and DNA methylation data across spatial replicates, our study provides critical insight into the effects of localized but strong fishing pressures on phenotypic change, informing conservation strategies, fisheries management and ntMPA design aimed at maintaining molecular biodiversity.

## Materials & Methods

### Study species and fishery

*X. novacula* (Labridae) is a protogynous hermaphrodite, with individuals first developing as females and transitioning into males later in life [29]. Adults occupy extensive sandy areas and defend adjacent small territories [30] (up to six female territories can be comprised within a larger territory of a male) forming polygynous social units [30,31]. During the reproductive period, fertilized eggs are released into the water column and are dispersed by currents [32]. After approximately 30 days, juveniles settle in coastal habitats [33], at locations distinct from their natal site. No adult migration has been observed, indicating strong site fidelity after settlement [34]. Together, these life-history traits imply limited local recruitment and a predominantly open population structure, where local adult populations are largely replenished by juveniles originating from multiple, hydrodynamically connected areas.

In the Balearic Islands, this species is heavily targeted by recreational fisheries and protective measures have been enforced such as a closed season during reproduction and bag limits [35]. However, the popularity and fishing pressure on this fishery has been increased significantly in recent years. It is estimated that in some locations up to 60% of the individuals can be harvested within just a few days following the opening of the fishing season [36]. Furthermore, fishing with natural bait (the type of capture that targets this species) selects for bigger, active and dominant individuals with larger territories (i.e. males). The effects of this localized but strong fishing pressure are currently under study but differences in density, individual sizes, ages and age at sex transitioning have been already detected between individuals inhabiting fisheries compared to the ones inhabiting ntMPAs [36,37]. The sedentary nature of the adults, combined with the dispersal capabilities of their eggs and larvae and the selective fishing pressure, make this species an ideal model to investigate life-history and molecular changes associated with trait-selective harvesting, as well as the role of ntMPAs in preserving molecular biodiversity.

### Sample collection and replicated experimental design

Individuals of *X. novacula* used in this study (N = 120) were captured in three different coastal areas (Palma, Llevant and Menorca) of the Balearic Islands (NW Mediterranean, Spain; Suppl. Fig. S1) in July-August 2022. Within each area, the samples were collected in two adjacent locations: a long-established ntMPA and a highly exploited fishery (Suppl. Fig. S1). For each location, we sampled 20 individuals to represent the range of age classes observed at the site, using the standard capture method for the species, baited hook-and-line gear. As source of genetic material, we collected a section of the pelvic fin (fin-clip) that was stored in ethanol 96º at 4ºC until DNA extraction. The otoliths were dissected and stored to obtain a direct measure of the individual’s age.

This study was positively evaluated by the Ethical Committee for Animal Experimentation of the University of the Balearic Islands (references for the protocols CEEA 224/10/23) and authorised by the Animal Research Ethical Committee of the Conselleria d’Agricultura, Pesca i Alimentació and the Direcció General de Pesca i Medi Marí of the Government of the Balearic Islands (reference for the authorisations SRM 34/2023 ERM and SSBA 04/2024 AEXP).

### Age determination

Age for each individual was estimated by studying the growth bands recorded in their otoliths following [38]. Briefly, otoliths were dissected and rinsed with hydrogen peroxide (4 %) and distilled water. The left otolith was then photographed under an optical microscope (Zeiss Axio Imager A1) and an AmScope MU900 USB2.0 eyepiece digital camera. Images were inspected for opaque and translucid bands alternating outward from the centre of the otolith. The succession of two different bands was considered one annual increment and its count an estimate for the age of the fish. This method has been successfully validated for *X. novacula* [39] and the readings were performed by an expert at the sclerochronology services of the Mediterranean Institute for Advanced Studies (IMEDEA).

### Phenotypic shifts under trait-selective harvesting

To evaluate phenotypic shifts resulting from trait-selective harvesting in our studied sample, we compared size and age distributions between protection levels. Differences in body size (cm) were assessed using a Linear Mixed Model (LMM) and age (count data in years) using a Generalized Linear Mixed Model (GLMM) with a Poisson distribution. In both models, protection level was included as a fixed effect and location as a random effect to account for spatial structure. Likelihood ratio tests were used to evaluate the significance of protection level by comparing models with and without the effect of protection. Analyses were conducted in R [40] v.4.2.3 using the *lme4* v.1.1.34 package [41].

### Laboratory analyses

DNA was extracted from fin-clips using the DNeasy blood and tissue kit (Qiagen N.V.) following the manufacturer’s protocol adding a RNase A (Qiagen N.V.) treatment. After extraction, DNA concentration was standardized by dilution in miliQ water to ensure equal contribution of all samples in the pooled genomic library.

In order to characterize genomic variation (single-nucleotide polymorphisms, SNPs) and DNA methylation levels, we used epi-genotyping by sequencing (epiGBS), a reduced-representation approach that combines enzymatic DNA fragmentation using the CpG-sensitive restriction enzyme *MspI* with bisulphite conversion. This method allows for the simultaneous assessment of DNA sequence and methylation variation in genomic regions enriched for CpG-dense sites, which in vertebrates often include gene regulatory regions. We implemented an adapted version of the laboratory protocol described in Gawehns et al. [42], optimised for sequencing on an Illumina NovaSeq 6000 sequencing platform. The detailed protocol of the new epiGBS3 methodology is provided as Supplementary Materials (Suppl. File epiGBS3 Protocol). Briefly, samples were digested with the restriction enzyme *MspI* (New England BioLabs Inc.) following the manufacturer’s protocol. Adapters with barcodes were ligated to the DNA digested samples using DNA ligase T4 (New England BioLabs Inc.). After adapter ligation, samples were pooled, cleaned and concentrated with the Nucleospin Gel & PCR Cleanup kit (MACHEREY-NAGEL GmbH & Co.) and a Nick repair treatment (New England BioLabs Inc.) was performed. The bisulphite conversion treatment was done using the DNA methylation lightning kit (ZYMO Research). The library was amplified by 15 cycles of the polymerase chain reaction (PCR) using the KAPA HIFI Uracil + hotstart ready mix (Roche Molecular Systems, Inc.). Finally, the library was cleaned and size-selected with Nucleospin Gel & PCR Cleanup kit and AMPupre XP beads (Beckman Coulter, Inc.). Individuals from different locations were randomized across the genomic libraries (i.e. all genomic libraries contained individuals from all locations). Custom made genomic libraries were sent to Novogene (UK Company Limited) for sequencing (150 bp pair end, PE).

### EpiGBS pipeline to extract genetic and DNA methylation data

The pipeline described in Gawehns et al. [42] was updated to improve efficiency and reproductivity and adapted to the new characteristics of the laboratory sequencing data. The new version, built on *Snakemake* [43] v7.1.0 and run within *Singularity* [44] v3.7.1 was used to process raw reads and obtain information on genetic variants and DNA methylation across samples. A full description of the bioinformatics workflow is provided in the Supplementary Materials (Suppl. File epiGBS3 Protocol). Briefly, raw multiplexed PE reads were filtered for polyG sequences using *fastp* [45] v.0.23.2 prior to loading into *Stacks* [46] v2.61 for PCR clone removal, sample demultiplexing and separation of positive and negative DNA strands. Adapters were trimmed using *cutadapt* [47] v4.1 and read quality was assessed before and after trimming using *Fastqc* [48] v0.11.4. PE clean reads were aligned to the reference genome assembly version fXyrNov1_1 [49] using the bisulphite-aware aligner *Bismark* [50] v0.24, which also provided cytosine methylation calls from the aligned reads.

### Statistics and bioinformatics for genetics data

#### - SNP calling and data filtering

To mitigate erroneous SNP calls introduced by bisulphite conversion, sorted alignment files were pre-processed using a double masking procedure that identifies non-complementary base pairs between DNA strands and masks bases that may represent bisulphite-induced artefacts (procedure included in the epiGBS3 bioinformatics pipeline; Suppl. File epiGBS3 Protocol). Masked alignment files for each sample were merged using *SAMtools* [51] v.1.7 with the *merge* function. The resulting file was used as input for variant calling with *FreeBayes* [52] v1.3.6. The output was a variant calling format (VCF) file, which was sequentially filtered using *Bcftools* [51] v1.13 and custom Python scripts. The filtering criteria were as follows: retaining only biallelic SNPs, removing calls with quality below 50, removing calls with sequencing depth below 7, removing high coverage loci within individual (loci with coverage above 99^th^ percentile), retaining loci present in a minimum of 10 individuals per location, excluding loci with a minor allele frequency (MAF) below 0.01 and pruning loci in linkage disequilibrium (LD) within a 20,000 bp window at an LD threshold above 0.5.

#### - Population genomics analyses

To assess genetic differentiation and structure between areas and protection levels, an Analysis of Molecular Variance (AMOVA) was conducted in R using the package *poppr* [53] v. 2.9.4. Statistical significance of variance components was assessed via 999 permutations. Location-level measures of nucleotide diversity (π) and pairwise F_*ST*_ values were estimated using the *populations* module [54] in *Stacks* v.2.68, with the filtered VCF file as input.

To evaluate differences in nucleotide diversity between protection levels, we fitted a LMM in R using the package *lme4*. SNP-level nucleotide diversity (π) per location was modelled as the response variable, with protection level and geographical area included as fixed effects. Area was treated as a fixed effect to reflect the block design and control for spatial heterogeneity. SNP identity was included as a random effect to account for locus-specific variability in diversity. The *bobyqua* optimizer was used to facilitate model convergence. The statistical significance of the fixed explanatory variable protection level was assessed by a likelihood ratio test.

To identify individual SNPs exhibiting significant differences in allele frequencies between protection levels, genotypic data was recorded as allele counts, representing the number of alternative and reference alleles per individual. For each SNP, allele counts were modelled using a binomial generalized linear model (GLM). Protection level and area were included as fixed effects. To control for multiple testing, p-values were adjusted using the *qvalue* function [55] v.2.30 in R.

Finally, to identify potential outlier loci suggestive of selection, we used the R package *pcadapt* [56] v.4. The number of principal components (K) was determined based on a scree plot of eigenvalues. Loci with significant Mahalanobis distances after multiple testing correction (q < 0.01), were considered candidates for selection.

### Statistics and bioinformatics for DNA methylation data

#### - DNA methylation data filtering and annotation

Bismark methylation calling reports were imported into R with the package *methylKit* [57] v1.16.1. After loading, DNA methylation data from CpG sites was destranded (merging of DNA methylation information from complementary CpG dinucleotides). Using custom R scripts adapted from Sepers et al. [58], DNA methylation data was filtered by the following criteria: removing sites by minimum sequencing depth of 10, removing high coverage sites within individual (sites with coverage above 99^th^ percentile), removing invariant sites across all individuals, removing fully methylated (methylation proportion > 0.95) and fully unmethylated (methylation proportion < 0.05) sites that are invariant in 85% of individuals and maintaining sites only if present in at least 6 individuals per location. Filtered data was used to obtain read counts of methylated and unmethylated cytosines and DNA methylation proportion (ratio of methylated to total reads) per individual and CpG site.

CpG sites were annotated to genomic features of the reference genome annotation for *X. novacula* [49] using the R package *GenomicRanges* [59] v.1.50.2. Promoter regions were defined as the regions 2000 bp upstream of the transcription start site (TSS) for genes on the positive strand and 2000 bp downstream for genes on the negative strand. In cases where a promoter overlapped with another annotated feature (e.g. exon or intron), the overlapping feature was assigned to prevent ambiguous classification.

Fishing reduces the abundance of older individuals, resulting in younger age structures in exploited populations [60]. Because DNA methylation at specific CpG sites can change predictably with age (epigenetic clock; [61,62]), differences between protected and exploited populations could partly reflect underlying age structure. Due to the inherent correlation between age and protection level, both variables cannot be included in the same statistical framework without violating independence and introducing collinearity. To address this, we used an age-reference subset of individuals from ntMPAs (displaying the full age range observed in the species), to identify age-associated CpG sites. For each CpG site, we fitted a Generalized Linear Model (GLM) in R, using counts of methylated and unmethylated reads as the response variable and including age and location as fixed effects. Models were fitted with a quasibinomial error structure to account for overdispersion and p-values were corrected for multiple testing using the *qvalue* package in R. All CpG sites significantly associated with age (q < 0.05) were removed from the dataset prior to downstream statistical analyses.

#### - DNA methylation statistical analyses

To evaluate DNA methylation differences across protection levels and geographical areas, a Permutational Multivariate Analysis of Variance (PERMANOVA) was performed in R using the package *vegan* [63] v.2.6-4. Manhattan distances were calculated between individuals based on CpG site methylation proportions and the resulting distance matrix was used in a PERMANOVA with the *adonis2* function. Statistical significance was assessed using 999 permutations.

To identify CpG sites exhibiting differential DNA methylation between protection levels, we implemented the same approach than described previously to detect CpG sites associated with age using GLMs. Methylated and unmethylated read counts were modelled as a two-column response using a quasibinomial error distribution. Protection level and area were included as fixed effects to test for the effect of protection and account for spatial structuring. P-values were adjusted to correct for multiple testing using the *qvalue* package in R. In addition to statistical significance, effect sizes were estimated using the package *emmeans* [64] v.1.10. A volcano plot for visual representation of the results was obtained using the R package *EnhancedVolcano* [65] v.1.16.0.

To assess genomic feature overrepresentation, we compared the genomic annotations of protection-level associated CpG sites (categorized as promoter, exon, intron and intergenic; and as coding vs. non-coding) to the expected distributions derived from the complete set of sequenced high-quality CpG sites. Fisher’s exact tests were used to evaluate deviation in annotation frequencies from the genomic background.

The differentially methylated sites (DMS) with large effect sizes (defined as absolute differences in DNA methylation proportion > 15%) were manually inspected based on gene annotations from the *X. novacula* reference genome [49].

## Results

### Data filtering and dataset composition

The initial variant calling performed with *FreeBayes* identified approximately 5 million variant calls relative to the reference genome, including low-confidence calls and sequencing artefacts. Following the application of the successive filters detailed in the Methods section, a total of 9,707 high-quality SNPs were retained for downstream analyses. The largest reduction in high-quality SNPs occurred during filtering for loci shared across individuals, a step designed to ensure robust comparison across locations.

The initial DNA methylation dataset generated using *Bismark* comprised approximately 2 million CpG sites. After applying filters, 49,344 high-quality CpG sites were retained for downstream analyses. As for the SNP dataset, the greatest reduction occurred when retaining only CpG sites present across individuals.

These substantial reductions in both SNPs and CpG sites when filtering for loci shared across individuals are inherent to reduced-representation sequencing protocols. Stochastic variation in library preparation and sequencing depth leads to heterogeneity in the subset of loci recovered across individuals. Therefore, the observed loss of loci reflects the nature of the methodology rather than a methodological artefact, compromised data quality or inadequate filtering strategies.

### Phenotypic shifts under trait-selective harvesting

Individuals collected from ntMPAs were significantly larger and older than those from exploited areas, indicating a clear demographic effect of harvesting in the sampled locations. On average, fish in ntMPAs were 2.5 cm longer than those in fisheries, with a mean total length of 17.4 cm and 14.9 cm respectively (N = 120, 95% CI = 0.3–4.6, χ^2^ = 4.66, p = 0.03; Suppl. Table S1). For age, individuals in ntMPAs were approximately 2.1 times older than those in fisheries, corresponding to an average age of 3.9 years and 1.9 years respectively (N = 100, 95% CI = 0.4–1.1, χ^2^ = 9.29, p = 0.002; Suppl. Table S1). These differences are consistent with the expected demographic effects of trait-selective harvesting, which removes from the population larger and older individuals.

### Population genomics analyses

The AMOVA results (Table 1) showed that 99.9% of the genetic variance was attributable to differences among individuals within locations. Geographic area and protection level nested within area explained less than 1% of the total variance, indicative of negligible genetic structuring across spatial or protection level factors. Consistent with these findings, pairwise F_*ST*_ values between locations calculated using *Stacks* were uniformly low (Suppl. Fig. S2), further confirming the absence of substantial genetic differentiation among locations.

**Table 1.** Results of the analysis of variance for the genetic (AMOVA) and DNA methylation (PERMANOVA) datasets. For each source of variation, the table reports degrees of freedom (df), the percentage of variance explained (Var) and associated p-values. P-values for both analyses were obtained using 999 permutations.

|  | Genetic AMOVA |  |  |  | DNA - Methylation PERMANOVA |  |  |
| --- | --- | --- | --- | --- | --- | --- | --- |
|  | df | Var (%) | p-value |  | df | Var (%) | p-value |
| Between Areas | 2 | - 0.03 | 0.58 |  | 2 | 2.35 | 0.05 |
| Between protection levels within areas | 3 | 0.07 | 0.09 |  | 3 | 3.59 | 0.03 |
| Within protection levels (residual) | 114 | 99.96 |  |  | 114 | 94.05 |  |
| Total | 119 | 100 |  |  | 119 | 100 |  |

Despite the lack of spatial genetic structuring, nucleotide diversity (π) significantly differed between geographical areas and protection levels. In terms of protection, ntMPAs exhibited significantly higher nucleotide diversity than their respective adjacent fisheries (χ^2^ = 22.43, p < 0.001; Fig. 1). Geographical areas also significantly influenced genetic diversity, suggesting spatial heterogeneity and a longitudinal gradient across the Balearic Islands (χ^2^ = 22.47, p < 0.001; Fig. 1; Suppl. Fig. S1)

**Figure 1.**
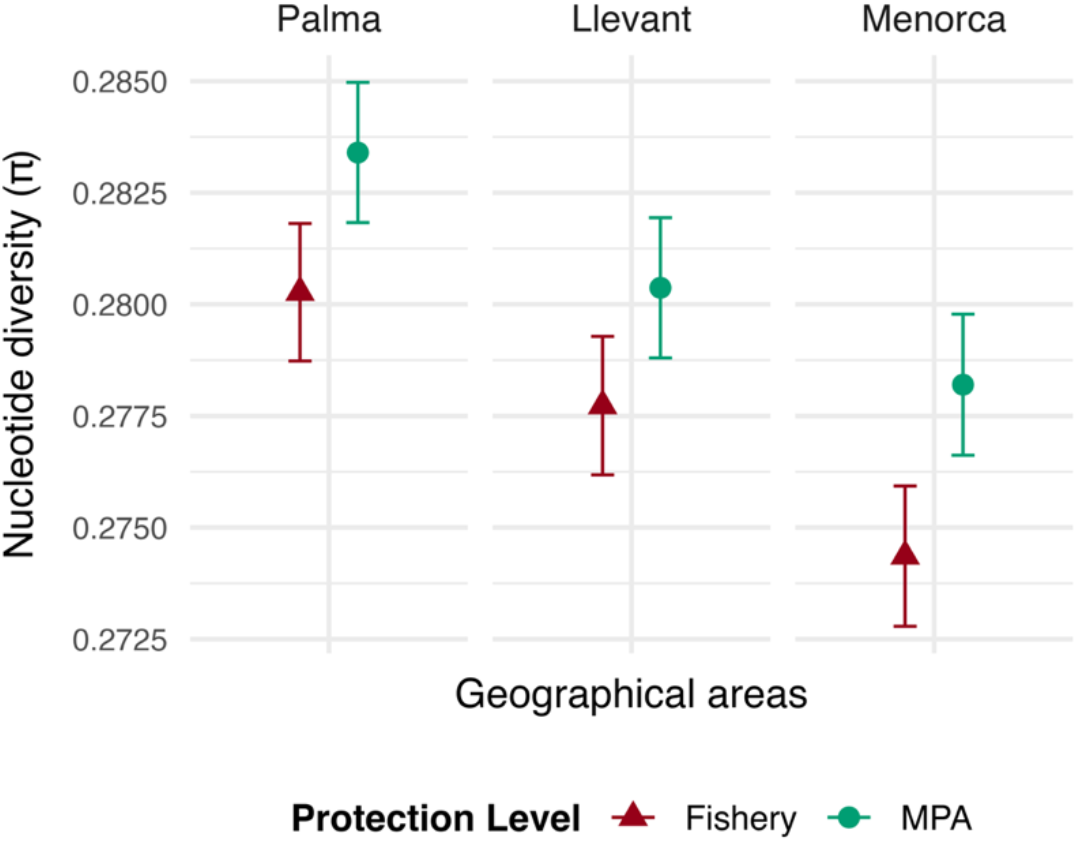
Nucleotide diversity (π) across the different study areas and protection levels.

To identify specific loci potentially associated with protection level, we implemented two approaches. The analysis of outlier loci using *pcadapt* detected no loci under selection at a significance threshold of q < 0.01, a result that aligns with the absence of genetic differentiation indicated by the AMOVA and F_*ST*_ analyses. We further examined potential differentiation between protection levels using GLMs at the SNP level. Of the tested SNPs, none remained statistically significant after multiple testing correction (q < 0.05). Overall, our results provide no evidence of genetic structure and selection associated with ntMPAs or fisheries at the genome-wide or locus-specific level.

### DNA methylation analyses

Genomic annotation revealed that the CpG sites result of quality filtering were distributed in similar proportions across introns, exons and intergenic regions (∼ 30 % each), and in smaller proportion (∼ 1 %) in promoter regions. Among genic CpG sites, approximately 90% were annotated to protein-coding genes. The low representation of promoters is expected, given that they constitute a minor fraction of the genome. In contrast, the similar proportions of CpG sites across exonic, intronic and intergenic regions, as well as the high representation of protein-coding genes, deviate from what would be expected under random DNA fragmentation. These patterns reflect the methodological bias of the epiGBS approach, which preferentially captures CpG-dense regions, such as promoters and gene bodies of protein-coding genes.

The PERMANOVA (Table 1) indicated that 94% of variance in DNA methylation was attributed to residual variation, while 2 % was explained by the area (p-value = 0.05) and 4% by protection level nested within area (p-value = 0.03). These results support the presence of significant differences in DNA methylation patterns between protection levels, after accounting for spatial structure and removing from the dataset age-associated sites.

From the GLM analyses, we identified a total of 291 significantly differentially methylated CpG sites (DMS; q < 0.05) between fisheries and ntMPAs (Fig. 2). The DMS deviated significantly in their genomic annotation profiles compared to the background CpG set. DMS were overrepresented in exonic and intronic regions, and underrepresented in intergenic regions. A modest, but statistically significant reduction was also observed in promoters. Additionally, DMS were significantly overrepresented in protein-coding regions (Table 2).

**Figure 2.**
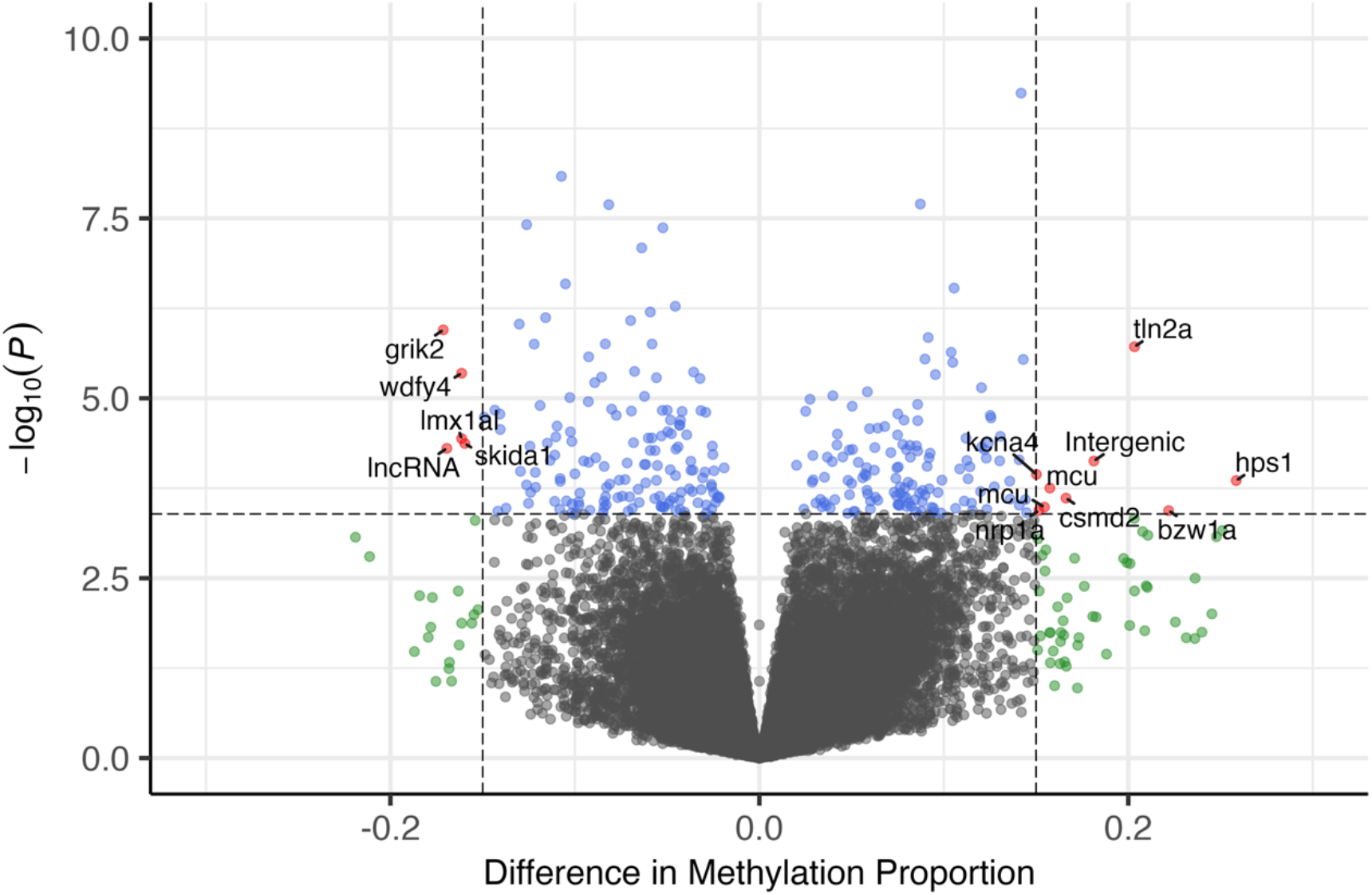
Volcano plot showing the distribution of CpG sites after removing age-associated sites. In red, sites that had a difference in DNA methylation between fisheries and no-take Marine Protected Areas larger than 15% and a q-value < 0.05. The labels correspond to gene annotation that includes the CpG site. One of the CpGs was not included within any known genetic boundary and was annotated as intergenic.

**Table 2.** Feature annotation differences for Differentially Methylated Sites (DMS) compared to the background dataset of all CpG sites included in the analysis. The upper section compares the differential distribution of annotated features and the lower section compares distributions among genic CpGs annotated to protein-coding versus non-coding genes. Statistical overrepresentation was evaluated using Fisher’s exact test. CI L = Lower Confidence Interval; CI U = Upper Confidence Interval.

|  | Percentage difference | Fisher's exact test |  |  |  |
| --- | --- | --- | --- | --- | --- |
|  |  | Odds ratio | CI L | CI U | p-value |
| Promoter | - 2.87 % | 0.54 | 0.28 | 0.96 | 0.03 |
| Exon | 8.30 % | 1.46 | 1.16 | 1.84 | 0.001 |
| Intron | 7.84 % | 1.41 | 1.12 | 1.77 | 0.001 |
| Intergenic | - 13.27 % | 0.50 | 0.38 | 0.66 | < 0.001 |
| Protein-coding<br>(in contrast to non-coding) | 6.85 % | 2.27 | 1.37 | 4.04 | < 0.001 |

A total of 14 DMS presented a DNA methylation level difference larger than 15% and were considered for further inspection (Fig. 2, Suppl. Table S2). Of those, 5 were hypermethylated in ntMPAs and 9 where hypermethylated in fisheries. The DMS hypermethylated in ntMPAs included a site in an exon of a long non-coding RNA (lncRNA; Xnov1ncA006550), a site in exon 1 of the *skida1* transcription factor and three sites located within introns of the genes *lmx1a* and *grik2* (associated with neural function) and *wdfy4* (involved in immune response). Among the DMS found hypermethylated in fisheries, one site was not annotated, likely located in an intergenic region, while remaining sites were located within introns of the following genes: *tln2a* (associated with cell signalling), *hps1* (melanin production), *mcu* (mitochondrial calcium import), *nrp1a* (vascular development), *csmd2* and *kcna4* (neural development and excitability) and *bzw1a* (protein synthesis).

## Discussion

Our results reveal that despite the absence of detectable genetic structure and no evidence of SNPs under selection, the fisheries studied here exhibited lower genetic diversity compared to adjacent ntMPAs. Furthermore, we found significant differences in DNA methylation among geographical areas and protection levels after removing age-associated CpG sites, suggesting that fishing pressure may influence epigenetic patterns beyond the known age-truncation effects. This provides the first empirical evidence that fisheries can induce epigenetic modifications via DNA methylation. Overall, this study contributes to a broader discussion on the role of ntMPAs in preserving genetic and epigenetic diversity of exploited fish populations.

Numerous studies have reported reduced genetic diversity in harvested populations [66,67], although the magnitude and ecological implications of these effects remain under debate and are underrepresented in management practices [68]. Genetic diversity loss is typically attributed to population declines that increase allele loss by genetic drift [69]. Sex- and size-selective harvesting further reduces effective population size (N*e*) and allelic richness [70]. In our study system, the lower genetic diversity in fisheries likely reflects such demographic effects. Groups of local demographic units of *X. novacula* function as a dynamic metapopulation, where local genetic diversity reflects the temporal accumulation of multiple cohorts originating from hydrodynamically connected source units rather than local recruitment alone. Under this framework, protected areas are expected to retain a greater number of cohorts and thus higher genetic diversity, whereas harvesting truncates age structure and reduces cohort representation even when census population size remains similar [36]. The observed reduction in nucleotide diversity in fisheries support this interpretation, while geographic variation among areas (Fig. 1) suggests that spatial heterogeneity can mask smaller, localized fishing effects. Together, these findings highlight the importance of considering age structure, cohort dynamics and spatial scale when assessing genetic impacts of exploitation and suggest that ntMPAs can act as reservoirs and sources of genetic diversity, particularly in open, dispersive systems.

DNA methylation patterns change predictably with age (epigenetic clock; [61,62]). Because selective harvesting truncates age structure, it is expected that DNA methylation patterns will be different between protected and harvested areas. To account for this, CpG sites associated with age were identified and removed, controlling for age-driven variation of DNA methylation. In sequential hermaphrodites, such as *X. novacula*, age, sex and body size are tightly linked, with older individuals being larger and predominantly male. Consequently, controlling for age also implicitly controls for sex- and size-associated DNA methylation effects. Despite this, DNA methylation patterns still differed consistently between protection levels, indicating that observed differences in DNA methylation cannot be explained by demographic structure alone.

*X. novacula* exhibits pelagic larval dispersal followed by settlement into either protected or exploited habitats, where individuals complete most of their growth and maturation. Thus, juveniles experience different post-settlement environments depending on protection status, which differ in population density, resource availability, predator abundance and social structures. Such different environmental conditions associated with harvesting could shape DNA methylation patterns in our study species, as multiple ecological and social factors, including population density, predation risk and social status, have been shown to influence DNA methylation patterns across taxa [71–75].

In this context, population density and the social environment experienced by juveniles of *X. novacula* during post-settlement ontogenetic stages may contribute to the DNA methylation differences observed between exploited and protected sites. Fishing reduces population density and selectively removes large, dominant individuals, potentially altering social hierarchies, mating opportunities and competitive interactions. Experimental studies demonstrate that early-life density and social conditions can shape DNA methylation patterns [72], and that dominance status is associated with distinct DNA methylation profiles in fishes [71]. In socially complex species, including *X. novacula* [31], selective removal of large, dominant individuals likely disrupts social structure in ways not fully captured by age alone. Beyond conspecific interactions, exposure to predators can induce DNA methylation changes [75]. As ntMPAs often harbour higher predator abundance (rays, sharks and dolphins), differential predation risk could further drive the observed DNA methylation differences between protection levels.

While DNA methylation is widely implicated in phenotypic plasticity, the extent to which DNA methylation patterns reflect environmentally induced responses, genetic background or their interaction is still under debate. Evidence suggests that gene expression regulation via DNA methylation can be shaped by underlying genetic variation [76]. In the context of fisheries, where environmental change and selective harvesting co-occur, disentangling these effects is particularly challenging. This study was not designed to separate genetic and environmental drivers of DNA methylation. However, the consistent divergence in DNA methylation between ntMPAs and fisheries, despite the absence of genetic structure, suggests an environmentally responsive component. At the same time, the absence of detectable genetic divergence does not rule out a genetic basis for DNA methylation differences, which may remain as undetected variation at loci influencing DNA methylation states.

Regardless of the causes and mechanisms driving the observed divergence in DNA methylation, the overrepresentation of differentially methylated sites (DMS) in gene bodies, particularly introns and exons, of protein-coding genes suggests potential regulatory effects at both transcriptional and post-transcriptional levels. Manual inspection of gene annotations at DMS revealed diverse potential functions, including neural development, immune response and cellular signalling. These annotations should be interpreted cautiously. Most are derived from targeted studies on model organisms under laboratory conditions and may not reflect gene function in wild marine fish exposed to complex environmental variability. Moreover, DNA methylation is highly tissue specific. In this study DNA methylation was assessed in fin tissue, associations with traits such as neural development are speculative, as gene function may vary across tissues and contexts. Although correlations between DNA methylation in peripheral tissues (e.g. blood) and other organs have been reported [77], the extent to which DNA methylation patterns and their functional consequences are shared across tissues remains unclear [77,78]. Future studies should validate the concordance of DNA methylation between dermal tissue and other target tissues, and clarify links with gene expression and splicing in *X. novacula*.

The use of ntMPAs and adjacent fisheries as a case-control design enabled comparisons of genetic and DNA methylation responses under contrasting selection regimes, while accounting for spatial structure and demographic complexity. However, the limited sample size may have prevented detection of SNPs under weak selection and DNA methylation differences with small effect sizes. Moreover, reduced representation sequencing with strict filtering likely excluded functionally relevant genomic regions. Our results also suggest that sampled locations belong to a highly connected metapopulation. To evaluate the true buffering capacity of ntMPAs, future work should assess the connectivity beyond the Balearic Islands and include populations with no history of exploitation to establish a true molecular baseline. Despite these limitations, our findings support the hypothesis that fishing-induced demographic and social disruptions may induce rapid, plastic responses mediated by DNA methylation. The identified DMS may contribute to local adaptation, functioning as a rapid and reversible mechanism of phenotypic variation in high gene flow marine systems.

In conclusion, we provide evidence that fisheries can induce epigenetic change in wild populations. From a management perspective, our findings highlight the importance of ntMPAs in preserving both genetic and epigenetic diversity, and the importance of understanding hydrodynamic connectivity to identify effective genetic reservoirs. A broader understanding of *X. novacula*’s population structure across its full distribution is essential for developing conservation strategies directed to preserve its molecular biodiversity.

## Acknowledgements

This work would have not been possible without the help of the IMEDEA Fish Ecology Lab team during field work, particularly Martina Martorell, Arancha Lana, Marco Signaroli and Eneko Aspillaga, as well as the staff at IEO-La Mola Field Station specially Juancho Movilla. We are also grateful to Raül Escandell for sharing his information on fisheries in Menorca and Oscar Gordo for his help in fieldwork and sample processing. Laboratory training at NIOO was generously provided by Martijn van der Sluijs.

