## Supplementary Information for "Fishing pressure induces changes in DNA methylation in genetically homogeneous marine metapopulations"

#### Supplementary figures

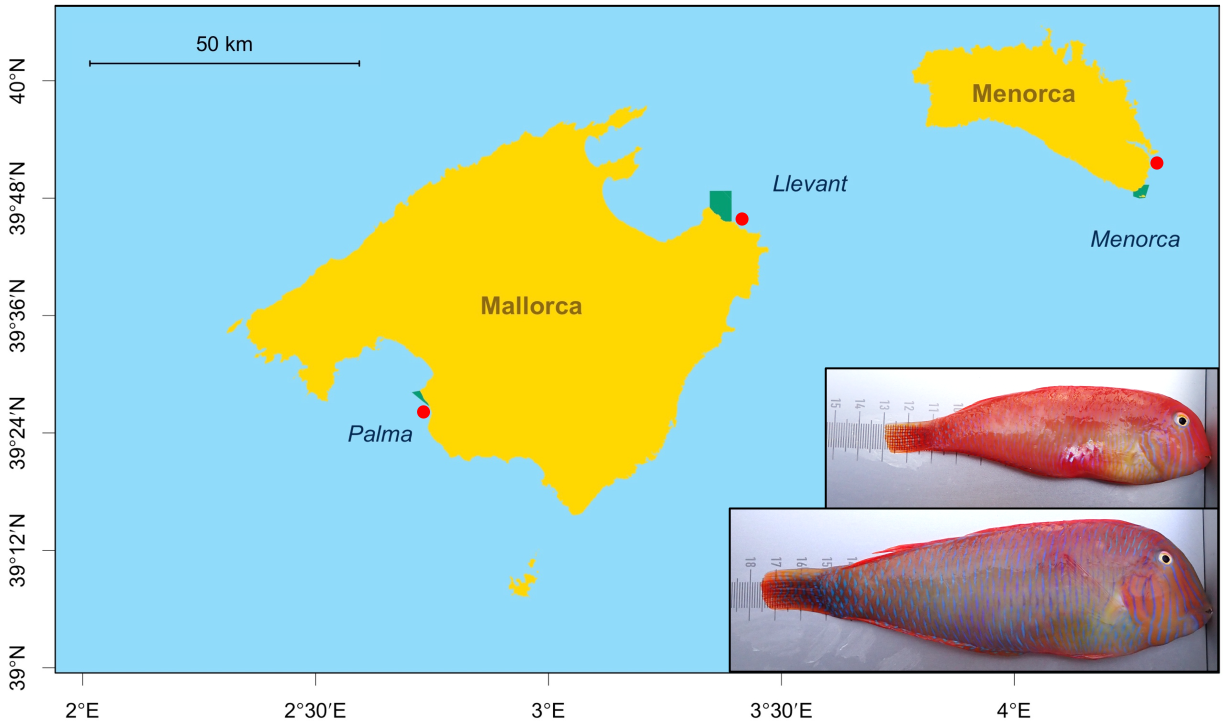


Supplementary Figure S1. Map of the Balearic Islands (Western Mediterranean Sea, Spain) showing the three sampled geographical areas (Palma, Llevant and Menorca). Within each area, two locations were defined: a fishery (Ses Olles, Cala Mesquida and Es Clot, respectively) in a red circle and a no-take Marine Protected Area (MPA Palma, MPA Llevant, MPA Illa de l'Aire, respectively) delimited by a green polygon depicting the area of legal protection. Female (top) and male (bottom) individuals of *Xyrichtys novacula*. Map created in R v.4.2.3 using the package *BTNtools* (https://github.com/aspillaga/BTNtools).

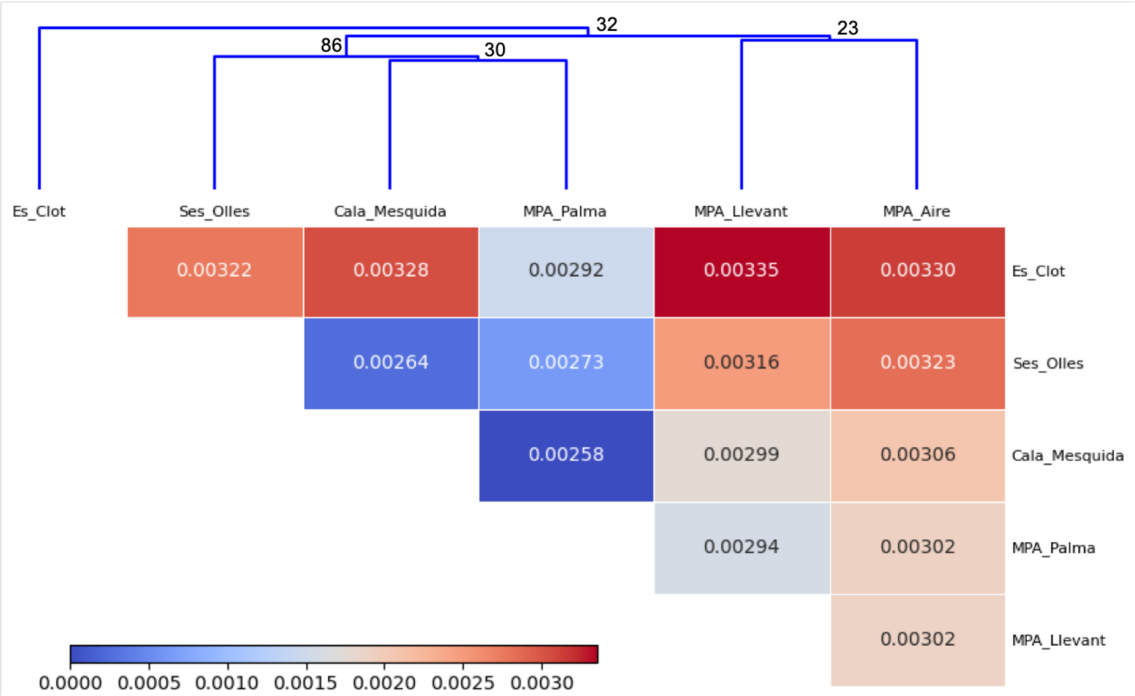


Supplementary Figure S2. Heatmap and dendrogram showing pairwise $F_{ST}$ estimates of genetic differentiation among sampling locations. The overall low values and weak bootstrap support for nodes (numbers on dendrogram) suggest limited genetic differentiation and a lack of clear population structure.

#### Supplementary tables

| | Model | Effect of ntMPA (Estimate) | 95% CI - L | 95% CI - U | p-value |
|---|---|---|---|---|---|
| Size (cm) | LMM | 2.46 | 0.31 | 4.60 | 0.031 |
| Age (years) | GLMM (Poisson) | 2.08 | 0.38 | 1.08 | 0.002 |

Supplementary Table S1. Results of the Linear Mixed Model (LMM) for body size (cm) and the Generalized Linear Mixed Model (GLMM) with a Poisson distribution for age (years), comparing individuals from fisheries (N = 60) and no-take Marine Protected Areas (ntMPA; N = 60). Estimates represent fixed effect coefficients; the estimate for age has been transformed to return to the original scale for interpretability. Statistical significance has been assessed using likelihood ratio test comparing the full model with the null model, without protection level. CI U = Upper 95% Confidence Interval; CI L = Lower 95% Confidence Interval.

| Chr. | Pos. | Effect | Feature | Loc. | Genome annotation ID | Gene | Gene name |
|---|---|---|---|---|---|---|---|
| 1 | 17360101 | Hyper Fish | Intron | Between exons 21-22 | Xnov1A031587 | tln2a | talin-2a |
| 8 | 15004806 | Hyper MPA | Intron | Between exons 3-4 | Xnov1A000215 | grik2 | glutamate receptor ionotropic, kainate 2 |
| 11 | 28567738 | Hyper MPA | Intron | Between exons 50-51 | Xnov1A020600 | wdfy4 | WD repeat- and FYVE domain-containing protein 4 |
| 11 | 29435419 | Hyper Fish | Intron | Between exons 1-2 | Xnov1A018983 | HPS1 | hybrid polyketide synthetase |
| 11 | 33039100 | Hyper Fish | Intron | Between exons 2-3 | Xnov1A018447 | mcu | calcium uniporter protein, mitochondrial |
| 11 | 33039116 | Hyper Fish | Intron | Between exons 2-3 | Xnov1A018447 | mcu | calcium uniporter protein, mitochondrial |
| 16 | 4431877 | Hyper MPA | Intron | Between exons 3-4 | Xnov1A019545 | lmx1al | LIM homeobox transcription factor 1-alpha-like |
| 18 | 21096145 | Hyper Fish | Intergenic | NA | NA | NA | Intergenic |
| 20 | 21571324 | Hyper MPA | Exon | Exon 1 (monoexonic) | Xnov1A037325 | skida1 | SKI/DACH domain-containing protein 1 |

| | | | | | | | |
|---|---|---|---|---|---|---|---|
| 20 | 8609450 | Hyper Fish | Intron | Between exons 7-8 | Xnov1A036326 | nrp1a | neuropilin-1a-like |
| 21 | 17489160 | Hyper Fish | Intron | Between exons 3-4 | Xnov1A023180 | csmd2 | CUB and sushi domain-containing protein 2 |
| 21 | 7087744 | Hyper Fish | Intron | Between exons 1-2 | Xnov1A035663 | kcna4 | potassium voltage-gated channel subfamily A member 4-like |
| 23 | 10326781 | Hyper MPA | Non-Coding | Exon 1 (monoexonic) | Xnov1ncA006550 | lncRNA | Long non-coding RNA |
| 24 | 19146392 | Hyper Fish | Intron | Between exons 13-14 | Xnov1A011423 | bzw1a | basic leucine zipper and W2 domain-containing protein 1-A-like |

Supplementary Table S2. Description of the CpG sites that presented a DNA methylation difference larger than 15% between protection levels and remained statistically significant after multiple testing correction ($q < 0.05$). Chromosome (Chr); Position (Pos.); Direction of the effect (Effect) being hypermethylated in fisheries (Hyper Fish) or hypermethylated in no-take Marine Protected Areas (Hyper MPA); Annotated feature that contains the CpG site (Feature); Location of the CpG site within the gene (Loc.); Identifier of the gene according to the genome annotation of the assembly version fXyrNov1_1 (Genome Annotation ID); Gene symbol (Gene); Full gene name (Gene name).

### Supplementary file epiGBS3 protocol

#### Background

EpiGBS (Gawehns et al., 2022; Sepers et al., 2023; van Gurp et al., 2016) was developed as a cost-effective method to assess CpG methylation in genomes using restriction digestion and bisulfite conversion. Instead of assessing methylation patterns of the whole genome, the restriction digestion allows the selection of a repeatable subset of the genome for which methylation is determined.

Here we present an adjusted wet-laboratory protocol and bioinformatics pipeline. We adapted the epiGBS2 protocol (Gawehns et al., 2022) to create sequencing libraries with dual indexing barcodes using TruSeq and Nextera indexing primers to permit sequencing on shared lanes on Illumina NovaSeq sequencers. The bioinformatics pipeline was adjusted to accommodate these changes during the demultiplexing steps. In addition to those, changes were made to deal with polyG reads resulting from two-colour base calling. The pipeline was also adapted to run with Singularity containers instead of Conda to facilitate reproducibility and overcome software library incompatibilities. Currently these changes of the bioinformatics pipeline have only been implemented for the reference-based version. A reference-free version will be added.

#### Modifications to the laboratory protocol

This laboratory protocol uses the laboratory protocol described in (Sepers et al., 2023) for great tit, *Parus major*, as a reference. It can be used for other species and has been tested for the species described in Table 1, with species-specific adjustments. Modifications have been made based on target number of reads, expected fragment size distribution and laboratory experience.

| Species | Samples per library | Enzymes | Specific changes compared to Sepers *et al.* 2023 |
|---|---|---|---|
| Great tit<br>*Parus major* | 24 | MspI - MspI | |
| Black grouse<br>*Lyrurus tetrix* | 24 | MspI - MspI | |
| Antarctic fur seal<br>*Arctocephalus gazella* | 24 | MspI - MspI | No beads before adapter ligation (step 4) but after step 7 |
| **Pearly razorfish**<br>***Xyrichtys novacula*** | **22** | **MspI - MspI** | **No beads before adapter ligation (step 4) but after step 7** |

Table 1. Species-specific adjustments to the laboratory protocol from Sepers et al., (2023). In bold the adjustments for the current study.

**Modification for Sequencing on NovaSeq 6000 Illumina platform (step 15 of the protocol)**

The home-made libraries were sequenced at Novogene (Novogene Europe, Munich) on a Illumina Novaseq 6000. Paired-end 150 bp, generating 3.3Gb raw data per sample, based on 22 samples per library (75Gb). Primers and adapters are provided in Table 2.

**Modification protocol annealing adapters (only first time)**

- Mix

  8 µl BA adapter I (10 µM) or CO adapter I (10 µM)
  8 µl BA adapter II (10 µM) or CO adapter II (10 µM)
  4 µl NEB buffer r2.1 (B6002S)
  20µl milliQ water
  ------------------------------------
  40 µl total (4 pmol/µl) = 4 µM

- Perform the annealing:

  5 min 95 °C
  Each minute -1 °C x 70 times (end temperature = 25 °C)
  5 min 4 °C

- Measure the concentration with the Qubit
- Dilute this annealed adapter mix in 5 mM Tris/HCl, pH 8.5 to 600 pg/µl.
- We verified the annealing on the Fragment Analyzer (Agilent, Middelburg, The Netherlands)
- Make a working stock by aliquoting the final dilution in portions
- Freeze all aliquots except the one in use, which can be stored for 2 months at 4 °C
- Avoid freezing-thawing cycles as much as possible!

A

| | | |
|---|---|---|
| BA-i5-adapter_I | 5’-AXAXTXTTTXXXTAXAXGAXGXTXTTXXGATXTZZZAAXTC-insert-CGGGAGGNNNGACAGAGAATATGTGTAGAGGCTCGGGTGCTCTG-3' | CO-i7-adapter_II |
| BA-i5-adapter_II | 3’-TGTGAGAAAGGGATGTGCTGCGAGAAGGCTAGANNNTTGAGGC-insert-CAGXXZZZXTGTXTXTTATAXAXATXTXXGAGXXXAXGAGAX-5' | CO-i7-adapter_I |

B

| | | | |
|---|---|---|---|
| GT primer A | F_i5-501-next | 5’-AATGATACGGCGACCACCGAGATCTACAC**TAGATCGC**ACACTCTTTCCCTACACGACGCTCTTCCGATCT-3’ | PAGE purified (IDT) |
| GT primer B | R_i7-701-next | 5’-CAAGCAGAAGACGGCATACGAGAT**TCGCCTTA**GTCTCGTGGGCTCGGAGATGTGTATAAGAGACAG-3’ | PAGE purified (IDT) |

Figure 1. EpiGBS3 adapters (A) and primers (B) with restriction sites translated to MspI. EpiGBS3 uses hemimethylated adapters. EpiGBS3 adapters consist of the Illumina NovaSeq adapter sequence (grey), a random 3 nucleotide sequence called UMI (yellow), a sample-specific barcode (red) and the complement restriction enzyme site overhang sequence (green), which includes a control nucleotide (blue). The 5′–3′ strand of the BA adapter and the 3′–5′ strand of the CO adapter contain only methylated cytosines (X); the opposite strands are unmethylated. All strands are dephosphorylated, therefore only adapter 3′ ends and DNA fragment 5′ ends ligate. This process generates single-strand nicks in the complementary strand, which are repaired during nick-translation using 5mC-dNTPs, resulting in fully methylated adapters. GT primers A and B contain the sequence that binds to the flow cell, the indexes (bold font) and the Illumina adapter sequences.

**Modifications to the bioinformatics pipeline**

The bioinformatics pipeline for the reference-based application has been adapted mostly for efficiency and repeatability. Other changes have been necessary to accommodate Illumina NovaSeq sequencing data.

The whole pipeline has been converted to run with publicly available Docker and Singularity containers instead of local Conda environments. This should make it easier to run the pipeline in different environments. To facilitate this, the contents of the python script that run clone_filter and process_radtags from Stacks (Rochette et al., 2019) have been moved to snakemake rules. To account for the dual indexing barcodes required for NovaSeq, the cutadapt (Martin, 2011) adapter trimming commands have been adjusted accordingly.

Other modifications are:

- Filtering of reads consisting of only polyGs using fastp (Chen, 2023). These polyG reads are the result of problems when sequencing the reverse read where the indexing primer is read correctly, but the sequencing read is not. This appears to be an artefact of the library design.
- Superfluous fields have been removed from the config file as they have no current use in the pipeline.

The code for the epiGBS3 pipeline is available on github.com/nioo-knaw/epigbs3.

| Company | Product | Purification | SequenceName | Barcode | Sequence |
|---|---|---|---|---|---|
| IDT | 100nmoleDNAOligo | PAGEPurification | GTprimerAF_i5-501-next | TAGATCGC | AATGATACGGCGACCACCGAGATCTACACTAGATCGCACACTC TTTCCCTACACGACGCTCTTCCGATCT |
| IDT | 100nmoleDNAOligo | PAGEPurification | GTprimerBR_i7-701-next | TAAGGCGA | CAAGCAGAAGACGGCATACGAGATTCGCCTTAGTCTCGTGGGC TCGGAGATGTGTATAAGAGACAG |
| IDT | 25nmoleDNAOligo | StandardDesalting | MspI_CO_II_WOB_2_CCAG_next | CCAG | CGGCTGGNNNCTGTCTCTTATACACATCTCCGAGCCCACGAGAC |
| IDT | 25nmoleDNAOligo | StandardDesalting | MspI_CO_II_WOB_3_TTGA_next | TTGA | CGGTCAANNNCTGTCTCTTATACACATCTCCGAGCCCACGAGAC |
| AlphaDNA | 200nmole | HPLC | MspI_CO_I_WOB_3_TTGA_next | TTGA | GTXTXGTGGGXTXGGAGATGTGTATAAGAGAXAGZZZTTGAC |
| AlphaDNA | 200nmole | HPLC | MspI_CO_I_WOB_2_CCAG_next | CCAG | GTXTXGTGGGXTXGGAGATGTGTATAAGAGAXAGZZZXXAGC |
| AlphaDNA | 200nmole | PAGEPurification | MspI_CO_I_WOB_5_ACTA_next | ACTA | GTXTXGTGGGXTXGGAGATGTGTATAAGAGAXAGZZZAXTAC |
| IDT | 100nmoleDNAOligo | StandardDesalting | MspI_CO_II_WOB_5_ACTA_next | ACTA | CGGTAGTNNNCTGTCTCTTATACACATCTCCGAGCCCACGAGAC |
| AlphaDNA | 200nmole | HPLC | MspI_BA_I_WOB_1_AACT | AACT | AXAXTXTTTXXXTAXAXGAXGXTXTTXXGATXTZZZAAXTC |
| AlphaDNA | 200nmole | HPLC | MspI_BA_I_WOB_2_CCTA | CCTA | AXAXTXTTTXXXTAXAXGAXGXTXTTXXGATXTZZZXXTAC |
| AlphaDNA | 200nmole | HPLC | MspI_BA_I_WOB_3_TTAC | TTAC | AXAXTXTTTXXXTAXAXGAXGXTXTTXXGATXTZZZTTAXC |
| AlphaDNA | 200nmole | HPLC | MspI_BA_I_WOB_4_AGGC | AGGC | AXAXTXTTTXXXTAXAXGAXGXTXTTXXGATXTZZZAGGXC |
| AlphaDNA | 200nmole | HPLC | MspI_BA_I_WOB_6_CCTTC | CCTTC | AXAXTXTTTXXXTAXAXGAXGXTXTTXXGATXTZZZXXTTXC |
| AlphaDNA | 200nmole | HPLC | MspI_BA_I_WOB_7_TTCAA | TTCAA | AXAXTXTTTXXXTAXAXGAXGXTXTTXXGATXTZZZTTXAAC |
| AlphaDNA | 200nmole | HPLC | MspI_BA_I_WOB_8_GCGGC | GCGGC | AXAXTXTTTXXXTAXAXGAXGXTXTTXXGATXTZZZGXGGXC |
| AlphaDNA | 200nmole | HPLC | MspI_BA_I_WOB_9_AGATGC | AGATGC | AXAXTXTTTXXXTAXAXGAXGXTXTTXXGATXTZZZAGATGXC |
| AlphaDNA | 200nmole | HPLC | MspI_BA_I_WOB_10_CATAGC | CATAGC | AXAXTXTTTXXXTAXAXGAXGXTXTTXXGATXTZZZXATAGXC |
| AlphaDNA | 200nmole | HPLC | MspI_BA_I_WOB_11_TTCGAC | TTCGAC | AXAXTXTTTXXXTAXAXGAXGXTXTTXXGATXTZZZTTXGAXC |
| AlphaDNA | 200nmole | HPLC | MspI_BA_I_WOB_12_ATGCGC | ATGCGC | AXAXTXTTTXXXTAXAXGAXGXTXTTXXGATXTZZZATGXGXC |
| AlphaDNA | 200nmole | HPLC | MspI_BA_II_WOB_1_AACT | AACT | CGGAGTTNNNAGATCGGAAGAGCGTCGTGTAGGGAAAGAGTGT |
| AlphaDNA | 200nmole | HPLC | MspI_BA_II_WOB_2_CCTA | CCTA | CGGTAGGNNNAGATCGGAAGAGCGTCGTGTAGGGAAAGAGTGT |
| AlphaDNA | 200nmole | HPLC | MspI_BA_II_WOB_3_TTAC | TTAC | CGGGTAANNNAGATCGGAAGAGCGTCGTGTAGGGAAAGAGTGT |
| AlphaDNA | 200nmole | HPLC | MspI_BA_II_WOB_4_AGGC | AGGC | CGGGCCTNNNAGATCGGAAGAGCGTCGTGTAGGGAAAGAGTGT |
| AlphaDNA | 200nmole | HPLC | MspI_BA_II_WOB_6_CCTTC | CCTTC | CGGGAAGGNNNAGATCGGAAGAGCGTCGTGTAGGGAAAGAGTGT |
| AlphaDNA | 200nmole | HPLC | MspI_BA_II_WOB_7_TTCAA | TTCAA | CGGTTGAANNNAGATCGGAAGAGCGTCGTGTAGGGAAAGAGTGT |
| AlphaDNA | 200nmole | HPLC | MspI_BA_II_WOB_8_GCGGC | GCGGC | CGGGCCGCNNNAGATCGGAAGAGCGTCGTGTAGGGAAAGAGTGT |
| AlphaDNA | 200nmole | HPLC | MspI_BA_II_WOB_9_AGATGC | AGATGC | CGGGCATCTNNNAGATCGGAAGAGCGTCGTGTAGGGAAAGAGTGT |
| AlphaDNA | 200nmole | HPLC | MspI_BA_II_WOB_10_CATAGC | CATAGC | CGGGCTATGNNNAGATCGGAAGAGCGTCGTGTAGGGAAAGAGTGT |
| AlphaDNA | 200nmole | HPLC | MspI_BA_II_WOB_11_TTCGAC | TTCGAC | CGGGTCGAANNNAGATCGGAAGAGCGTCGTGTAGGGAAAGAGTGT |
| AlphaDNA | 200nmole | HPLC | MspI_BA_II_WOB_12_ATGCGC | ATGCGC | CGGGCGCATNNNAGATCGGAAGAGCGTCGTGTAGGGAAAGAGTGT |

Table 2. Sequence information for the primers and barcodes used for this study. Given are the company name (IDT = Integrated DNA Technologies, Leuven, Belgium; AlphaDNA, Montreal, Canada) where the product was acquired, the purification method, the sequence name, the barcode and the DNA sequence. The Barcode of the GTprimerBR_i7-701-next adapter, is the reverse complementary with the original one. Sequences are: degenerate N = [ACTG], Z=[ATXG] X=[5-Me-dC].
